# macpie: scalable workflow for high-throughput transcriptomic profiling

**DOI:** 10.1101/2025.08.06.669002

**Authors:** Nenad Bartonicek, Xin Liu, Laura Twomey, Michelle Meier, Richard Lupat, Stuart Craig, David Yoannidis, Jason Li, Tim Semple, Kaylene J Simpson, Mark X Li, Susanne Ramm

## Abstract

High-throughput transcriptomic profiling (HTTr) enables scalable characterisation of transcriptional responses to chemical and genetic perturbations. While plate-based technologies such as MAC-Seq, TempO-seq and PLATE-seq have made HTTr more accessible, they pose unique computational challenges in modelling data and integration across modalities. We present *macpie*, an R package designed to streamline the analysis of HTTr data from plate-based screens. Built on the tidySeurat framework, *macpie* streamlines the entire analytical pipeline from preprocessing and quality control to pathway enrichment, chemical feature extraction, and multimodal data integration. The package incorporates multiple statistical frameworks and leverages parallelisation for scalability. By leveraging Docker and Nextflow, macpie ensures reproducibility and ease of use for transcriptome-wide screening.

**Availability:** The R package *macpie* is freely available at https://github.com/PMCC-BioinformaticsCore/macpie, with images of the working environment hosted at Docker Hub: xliu81/macpie. A companion Nextflow pipeline for preprocessing from FASTQ files is available at https://github.com/PMCC-BioinformaticsCore/dinoflow.

**Supplementary information:** Package vignettes with the full analytical workflow available at *https://pmcc-bioinformaticscore.github.io/macpie/articles/macpie.html*

## Introduction

High-throughput screening (HTS) platforms are widely used in modern biomedical research, supporting a broad range of applications from functional genomics (e.g., CRISPR or RNAi screens), biomarker identification, drug discovery, to toxicology^1-4^. Early HTS approaches commonly reduced complex cellular phenotypes to a limited set of features such as proliferation rates, morphological changes or biomarker abundance, and were gradually replaced with multi-feature, high-content screening (HCS)^5^. More recently, high-throughput transcriptomic (HTTr) profiling has emerged as an increasingly accessible and scalable extension to HCS, capturing dynamic cellular states in response to chemical or genetic perturbations^6-8^.

Several technologies support HTTr, including DRUG-seq^9^, TempO-seq^10^, L1000^6^, PLATE-seq^11^, and commercial platforms such as Insphero’s Organoid DRUG-seq^12^. MAC-Seq (Multiplexed Analysis of Cells) was introduced as a cost-effective HTTr method that eliminates RNA extraction steps and supports integration with suspension cells, flow cytometry read outs and high-content imaging for adherent, 3D matrix embedded cell models, as well as complex co-culture systems^13^. Even though these platforms overcome considerable technical barriers related to plate-based workflows, they also present unique analytical challenges.

High-throughput transcriptomic platforms generate datasets that fall outside the scope of existing computational workflows. Compared to conventional RNA-seq, HTTr datasets often involve complex experimental designs with larger number of perturbations and higher potential for latent batch effects. The limited input material and small cell numbers frequently result in zero-inflated count distributions, resembling those seen in single-cell RNA-seq (scRNA-seq)^14^. However, HTTr datasets generally lack the number of samples needed to support statistical models developed for scRNA-seq, presenting unique analytical challenges.

Current HTTr analytical frameworks were mostly built to address toxicology analyses and are available as standalone applications (*BMDExpress-2*^15^) or partial workflows in R for quality control (e.g. *httrpl*^8^), analysis and visualisation (e.g. DRUG-seq^16^) or modelling of compound concentration-response curves (e.g. *tcplfit2*^17^, and *bmd*^18^). A complete, modular, and scalable R-based framework integrating both bioinformatics and cheminformatics components remains absent. To address this gap, we introduce *macpie*, an R package for the comprehensive analysis of HTTr data from plate-based sequencing technologies.

## Implementation

The *macpie* package was developed for analysis and visualisation of HTTr data, particularly for large, 384 well plate-based screens. The workflow is based on the *tidySeurat* framework that combines properties of *Seurat* and *tidyverse* objects, benefitting from their respective functionalities. *Seurat* is a widely used R package for complete analysis of transcriptomic data from single cell experiments^19^, including QC, dimensionality reduction, clustering, marker selection and data integration. The *tidyverse* collection of R packages is the standard in R-based data analytics^20^ for preprocessing, modelling and visualisation of complex data structures. *macpie* is compatible with R version >4.3.3 and is available as a Docker container to ensure reproducibility and ease of installation. Key steps of the analytical workflow are outlined in the package vignettes.

### Data preprocessing

*macpie* requires gene expression matrices and corresponding sample metadata sheet as inputs. Matrices can be imported from the standard sparse matrix format, as generated by Cell Ranger^22^ or STARsolo^23^ from raw FASTQ files. The sample metadata sheet must contain a minimal set of standard columns as outlined in the documentation, ensuring accurate mapping of expression values to sample annotation via sample barcodes, and can be further populated by descriptors of samples or perturbations. To simplify and standardise data preprocessing from FASTQ files, we provide a companion Nextflow pipeline, available at https://github.com/PMCC-BioinformaticsCore/dinoflow. *macpie* currently supports data from *Homo sapiens* and *Mus musculus*.

### Quality control

Given the large-scale nature of HTTr transcription-based screens, quality controls are important to ensure data integrity and minimise technical artefacts (Fig 1A). The QC workflow begins with metadata validation, as screens involve complex, often manually entered combinations of treatments. To streamline this step, we simplified inspection of large numbers of experimental variables for potential errors or inconsistencies, including automated checks for missing values and special characters. Next, *macpie* provides a set of QC tools that assess various properties of read distribution across genes and experimental conditions: read depth, variability, variance decomposition, outlier detection and normalisation. Relative log expression (RLE) plots are especially effective at visualising impacts of filtering cutoffs and various normalisations methods, while quantifying differences with the average coefficient of covariation (Fig 1A).

**Figure 1.**
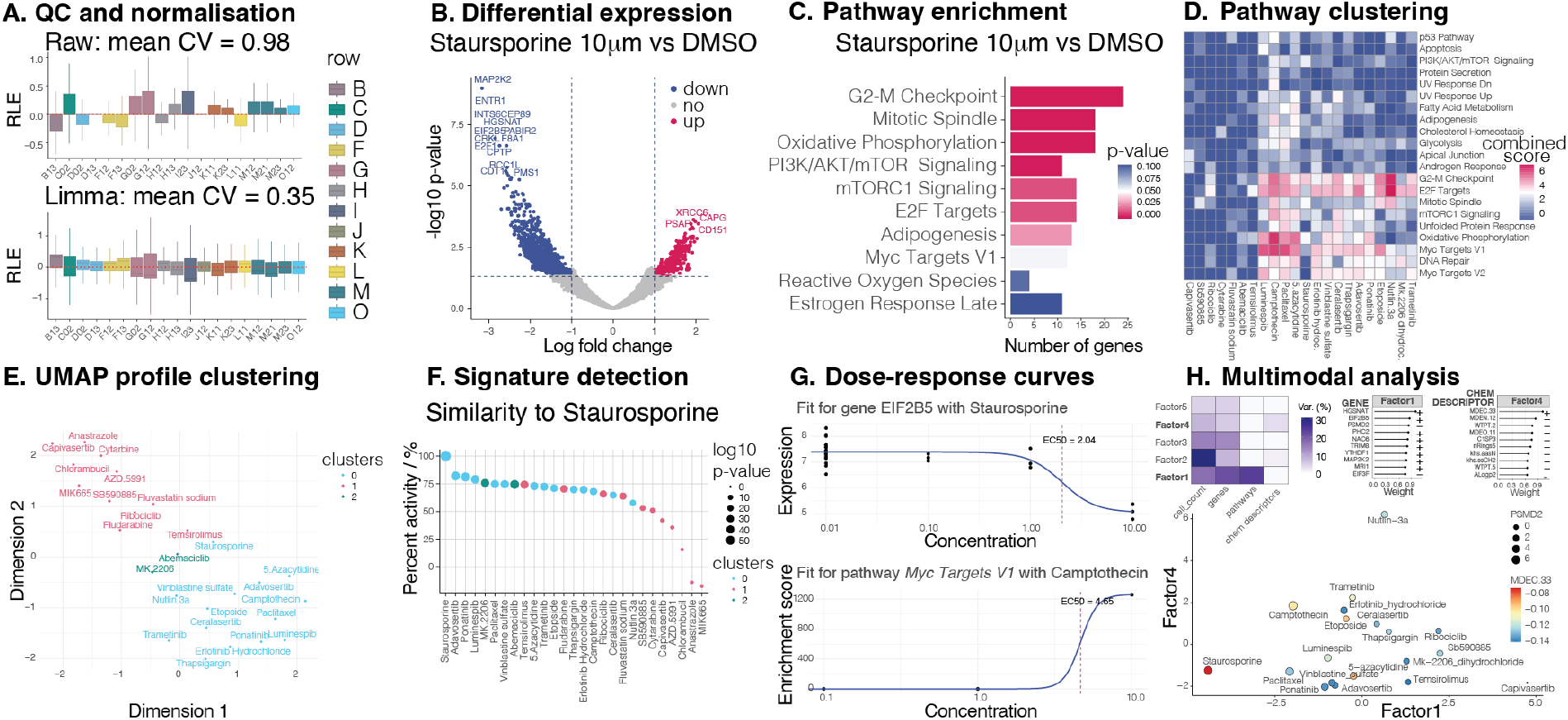
Overview of *macpie* key functionalities. A. Relative log expression (RLE) plots to estimate sources of variability during quality control and normalization. B. Volcano plot showing differential gene expression (DE) following Staurosporine 10 µM treatment vs DMSO vehicle control. C. Pathway enrichment heatmap derived from DE analysis. D. Aggregated pathway visualisation, combining pathway enrichment results across replicates. E. UMAP dimensionality reduction based on DE of treatments compared to DMSO. F. Point plot for signature detection, based on the enrichment of top 500 genes from an existing perturbation profile, as measured by Normalised Enrichment Score (NES). G. Dose-response curves at the gene level (top) and pathway level (bottom), with EC50 values indicated by dashed red lines. H. Multimodal analysis using MOFA^21^. Upper panels describe latent factors of a model that combines cell viability, gene expression, pathway enrichment and chemical properties of compounds. Lower panel shows low-dimensional representation of samples coloured by molecular descriptor MDEC.33, with data point size defined by expression of by PSMD2 gene, a cancer growth promoter.

### Transcriptomic workflow

Following QC, *macpie* supports two principal analytical modes: single-treatment and multi-treatment analysis. At the single-treatment level, *macpie* simplifies characterization of transcriptional responses to individual perturbations. Differential expression analysis is implemented with support for multiple statistical frameworks, including Seurat^19^, DESeq2^24^, edgeR^25^, RUVSeq^26^ and ZINB-WaVE^27^, allowing users to choose between models based on their dataset characteristics. Differentially expressed genes (DEGs) can be visualised on volcano plots (Fig. 1B), placed in biological context with gene set enrichment and pathway analysis tools (Fig. 1C), and samples visualised on multidimensional scaling (MDS) and Uniform Manifold Approximation and Projection (UMAP) plots. In addition, *macpie* offers a multi-treatment analysis framework, enabling users to compare transcriptional profiles across numerous perturbations in parallel. This mode automates the computation of DEGs and the enrichment analysis via standardized pipelines, producing summaries of results on gene and pathway levels (Fig D-F). All multi-treatment analyses are parallelized to optimize runtime and computational efficiency.

### Screen-level analyses

To explore higher-order effects across perturbations and integrate them with user-provided external annotations such as cell counts of surface markers, the *macpie* workflow streamlines several visualisations and analyses. First, *macpie* simplifies UMAP dimensionality reduction, enabling users to cluster experimental factors by the similarity of their transcriptional responses (Fig 1E). In addition, *macpie* supports cheminformatics workflows by extracting SMILES strings from compound names, computing and filtering molecular descriptors (e.g. physicochemical properties and fingerprints) or calculating half maximal effective concentrations for genes or pathways (Fig 1G). Finally, *macpie* allows data integration with Multi-Omics Factor Analysis (MOFA)^21^, facilitating unsupervised decomposition of variance across different data modalities, revealing latent factors driving biological or technical variation (Fig 1H).

## Conclusion

*macpie* offers a robust, well-structured and flexible R-based framework for the analysis of high-throughput transcriptomics screens. By standardizing preprocessing and integrating diverse statistical methods, it enables consistent and interpretable analysis of complex datasets. A complementary Nextflow pipeline for raw data preprocessing and containerized environment support were developed to provide scalability and reproducibility required for large-scale screening efforts. The modular architecture and emphasis on portability were designed to encourage community contributions to *macpie* in the future. We successfully applied *macpie* to other external HTTr datasets (vignette “cross_platform_compatibility”), confirming its robustness across different platforms, cell types, and perturbations.

As multi-modal profiling becomes standard for characterizing cellular responses, *macpie* provides a foundation for integrating transcriptomic and phenotypic data with chemical or biological properties of treatments. Future developments will incorporate additional omics layers, such as surface proteomics, secretomes, long-read sequencing and cellular barcoding^28^, facilitating more comprehensive systems-level analyses.

## Acknowledgements

We thank Ricky Johnstone, Magdalena Nakova and Hasan Quraishi for helpful discussions during the preparation of this manuscript, and Jenni Luu, Robert Vary, Kavya Pamulapati, and Karla Cowley from Victorian Centre for Functional Genomics (Peter Mac) for data generation. We acknowledge the Computational Biology Program at Peter MacCallum Cancer Centre and the Data Sprint initiative, including Miriam Yeung for her work on the Nextflow workflow. This work was supported by a Peter Mac Foundation Grant awarded to SR.

